# PW-GAN: Pseudo-Warping Field Guided GAN for Unsupervised Denoising of Fetal Brain MRI Images

**DOI:** 10.1101/2025.04.30.651598

**Authors:** Linxin Chen, Jiaying Lin, Guixing Fu, Pengjie Yang, Mengting Liu

**Affiliations:** School of Biomedical Engineering, Shenzhen Campus of Sun Yat-sen University, Shenzhen, 518107, China

**Keywords:** Fetal Brain MRI, Medical Image Denoising, Generative Adversarial Network, Unsupervised Learning, Pseudo-Warping Field

## Abstract

Fetal brain magnetic resonance imaging is of great importance for prenatal neural disorder diagnosis. To improve signal-to-noise ratio and coverage, fetal brain MRI often uses thick-slice scanning to reduce motion artifacts and ensure image quality for single slices. Reconstructing 3D MR volumes from multiple motion-corrupted stacks of 2D slices has shown promise in imaging of moving subjects like fetal MRI. However, raw scans acquired with varying slice thicknesses present substantial disparities in quality when retrospectively reconstructed into isotropic high-resolution volumes (e.g. 0.8 mm slice thickness). In particular, thick-slice acquisitions (e.g. 5-6 mm) tend to yield suboptimal reconstruction results compared to thin-slice scans (e.g. 2-3 mm), often exhibiting residual noise, interpolation-induced artifacts, and inter-slice structural discontinuities. This challenge has highlighted the need to further reduce noise caused by reconstruction errors when the reconstructed volumes are used for downstream tasks. With the success of neural networks in computer vision, deep learning methods based on architectures such as convolutional neural networks (CNNs) and generative adversarial networks (GANs) have achieved outstanding performance in single-image denoising. Nevertheless, CNN-based methods still heavily rely on large-scale datasets comprising noisy-clean image pairs. Although unsupervised GAN variants such as CycleGAN have been developed to mitigate this dependency, the inherent variability of GAN-generated outputs often results in tiny anatomical distortions, significantly limiting the applicability of GAN-based methods. While several approaches have introduced regularizations to address this issue, they largely focus on denoising individual slices, overlooking inter-slice structural inconsistencies that arise from treating slices independently. In this work, we propose Pseudo-Warping Field Guided Unsupervised Generative Adversarial Network (PW-GAN), which formulates post-reconstruction optimization as an unpaired style transfer problem between low-quality and high-quality MRI domains. Moreover, by incorporating a pseudo-deformation field module based on optical flow estimation, our method significantly enhances inter-slice continuity in the reconstructed volumes while effectively suppressing residual noise and interpolation artifacts introduced during the reconstruction process. Evaluations on both simulated and *in vivo* data demonstrate that our method outperforms existing unsupervised models and achieves performance on par with several state-of-the-art supervised methods.

## 1. Introduction

Fetal brain magnetic resonance imaging (MRI), as a high-resolution and non-invasive medical imaging technology, provides more comprehensive anatomical information and finer spatial details than ultrasound for prenatal diagnosis. It holds significant value in the evaluation of complex congenital brain abnormalities [1–4]. Nevertheless, the fetal brain contains numerous small and intricate anatomical structures and exhibits inherently low tissue contrast [5]. Moreover, image acquisition is highly susceptible to motion artifacts caused by spontaneous fetal movements and maternal respiration [6]. These challenges necessitate a careful balance in clinical MRI protocols among image signal-to-noise ratio (SNR), voxel size (and thus spatial resolution), and acquisition time. A commonly adopted clinical strategy is to maintain small voxel sizes in the phase and frequency encoding directions to suppress motion artifacts and achieve high in-plane resolution, while keeping the overall scan time and SNR within acceptable limits. This, however, comes at the cost of increased slice thickness and reduced through-plane resolution, resulting in anisotropic voxels [7].

Many super-resolution reconstruction (SRR) techniques [8–11] have been developed to integrate complementary information from multiple motion-corrupted stacks of 2D slices to reconstruct isotropic high-resolution 3D volumes from anisotropic acquisitions [12, 13]. However, the reconstructed image is still contaminated with noise, some of which arises from unidentified sources linked to the inherent nonlinearity of the reconstruction algorithm. This issue is particularly pronounced in reconstructions from thick-slice acquisitions (e.g., 5-6 mm) compared to those from thin-slice scans (e.g., 2-3 mm). The remaining noise might be attributable to multiple factors: First, thick-slice acquisitions are highly susceptible to partial volume effect (PVE), which blur the boundaries of fine brain structures [14, 15], this issue is difficult to resolve during the SRR process. Second, interpolation operations involved in SRR introduce non-physical intensity transitions at structural edges, manifesting as staircase-like discontinuities. These artifacts exhibit spatial correlations, fundamentally differing from the random distribution of acquisition noise, thereby rendering conventional denoising filters ineffective. Third, residual noise from the preprocessing stage undergoes nonlinear modulation during the SRR process, leading to an amplification of its impact. Critically, such reconstruction-induced artifacts and distortions lack the well-defined statistical characteristics like MRI system noise (e.g., Rician distribution) [16–18]. This challenge can be framed as a post-reconstruction image denoising and optimization problem, aiming to remove residual noise from the thick-slice SRR reconstructed images, and to improve the inter-slice consistency and fidelity of the resulting volumes [19]. For clarity, in the following sections, we will refer to the 3D volumes reconstructed from thick-slice data as noisy images and, analogously, term those from thin-slice reconstructions as clean images.

With the success of neural networks in computer vision, a number of deep learning based methods have been proposed for image denoising and optimization [20–23]. Similar methods are also applied to medical image denoising [24–27]. Initially, researchers trained convolutional neural network (CNN) with paired clean-noisy image pairs to learn a direct mapping from noisy image to its clean counterpart. However, CNN model typically adopt loss functions such as mean squared error (MSE) or mean absolute error (MAE), which tend to drive the network outputs toward the average of the noisy input. This often results in excessive smoothing, particularly degrading high-frequency details such as edges and textures. The advent of DnCNN [28], which integrates residual learning and batch normalization, has demonstrated impressive denoising performance and anatomical fidelity. Nevertheless, these approaches are predominantly supervised and rely heavily on large datasets with clean-noisy image pairs, a requirement that poses significant limitations for direct clinical practice.

To tackle this challenge, a number of unsupervised methods that rely solely on noisy images have also been explored over the past few years. Lehtinen et al. proposed Noise2Noise [29], which leverages pairs of noisy images with identical underlying content but differing noise realizations to train a model that statistically learns the expected clean images. Under the assumption of a known distribution of clean images, this approach enables denoising without the need for explicit clean-noisy pairs. Unfortunately, in clinical practice, acquiring such paired noisy images with consistent anatomical structures yet varying noise levels remains highly challenging. Building upon this idea, Noise2Void [30] and Noise2Self [31] further reduce dependency on clean data and introduce blind-spot strategies to infer clean signals from within the noisy observation itself. More recently, Deformed2Self [32] incorporates inter-slice motion cues by exploiting the spatial similarity and deformation between adjacent slices, thereby enhancing both denoising and image quality in volumetric data. Although the lack of clean reference images is an inherent limitation of unsupervised learning, this constraint significantly hampers the denoising performance of such models, making it difficult for them to surpass supervised approaches trained on paired clean and noisy images.

In recent years, generative adversarial network (GAN) have gained popularity in natural image denoising due to their ability to model global image distributions and capture perceptually relevant differences [23, 33]. By reformulating the denoising and enhancement task as a style transfer problem between the noisy and clean domains [34–36], GAN model is able to learn the distribution of clean images under unsupervised conditions. Yet, in the context of medical imaging, the emphasis on overall structural plausibility can lead to distortion of anatomical details—an outcome that is unacceptable in clinical settings. To tackle this challenge, several newer-generation GAN frameworks such as Conditional GAN (CGAN) [26, 37] and two-stage framework of GAN-CNN [25] have attempted to incorporate conditional information with the aim of preserving fine anatomical structures. However, the use of paired clean-noisy images causes these models to degrade into a supervised learning form. On the other hand, OTE-GAN seeks to prevent anatomical structure distortion by incorporating optimal transport theory. Despite their innovations, most of these models are primarily tailored for single-slice denoising, which makes it challenging to fully address the inter-slice discontinuities frequently encountered in fetal brain MRI reconstruction. To tackle this issue, one straightforward solution is to employ 3D image denoising models. However, the computational burden associated with 3D models remains prohibitively high, limiting their practicality and widespread adoption. An alternative and more feasible strategy is to introduce additional inter-slice regularization within 2D models. This can be achieved by incorporating pseudo-warping field [19, 39], where a random warping field is generated for each slice. Such an approach enhances the model’s sensitivity to fine anatomical structures and inter-slice discontinuities, thereby improving slice-to-slice consistency and the preservation of anatomical details without substantially increasing computational overhead.

In this work, we propose a deep learning framework based on GAN for post-reconstruction image denoising and optimization, enabling thick-slice fetal brain MRI reconstructions to be refined to a quality comparable with that of thin-slice results. To address the issue of inter-slice discontinuity and suppress interpolation artifacts introduced during reconstruction, we incorporate a pseudo-warping field. This field guides the model to improve spatial coherence across slices while denoising. We evaluated our model on reconstructed fetal brain MRI volumes using both simulated noise and real-world clinical datasets. The simulated noises are derived from a mathematical model to mimic the MRI reconstruction process and apply this to the Atlas CRL dataset [40]. This allows for direct qualitative and quantitative comparisons with supervised methods. The results show that our model achieves denoising performance on par with supervised baselines, while maintaining fine anatomical structures. Additionally, the results of ablation studies show that the introduction of the pseudo-deformation field significantly improves inter-slice continuity in thick-slice reconstructions.

In summary, the main contributions of this work are as follows:

1. We propose a PW-GAN-based post-reconstruction denoising and optimization framework tailored for thick-slice fetal brain MRI. By introducing a pseudo-warping field, our model promotes inter-slice continuity while eliminating the dependence for paired clean-noisy training data.
2. To preserve anatomical integrity, we incorporate multiple loss terms—cycle consistency, reconstruction consistency, and spatial consistency—that prevent anatomical distortion during denoising.
3. To further enhance the perceptual quality of edge and texture regions associated with anatomical detail, we employ a perceptual loss derived from a pre-trained VGG network [41], which strengthens the model’s capacity for high-level feature extraction.

## 2. Methods

### 2.1. Problem Formulation

Let 𝒳 and 𝒴 denote the reconstructed volumes from thick- and thin-slice inputs, representing the noisy and clean domains, respectively. The reconstruction process that generates isotropic high-resolution volumes is denoted by ℛ,

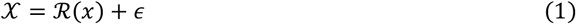

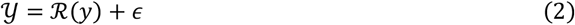

where *x* refers to the original thick-slice data before reconstruction, and likewise, *y* refers to the original thin-slice data. Notably, *ϵ* denotes a small, potentially inherent systematic error associated with the reconstruction process.

In practice, the original thick-slice data *x* typically contains residual noise that is difficult to eliminate completely. This can be expressed as:

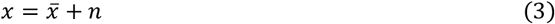

where 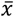 denotes the ideal, noise-free thick-slice data, and *n* represents the residual noise that remains after initial denoising.

Substituting Equation (3) into Equation (1), we obtain:

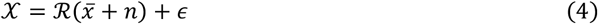

Furthermore, by separately considering the reconstruction process applied to each component and their interactions, Equation (1) can be expanded as:

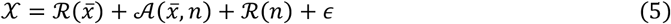

Here, ℛ(*n*) reflects how residual noise may be modulated or amplified during reconstruction, while 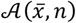 denotes a nonlinear interaction term capturing artifact formation arising from the interplay between anatomical structures and noise. For example, in the presence of noise, anatomical edges may become ambiguous, leading to reconstruction-induced distortions. Notably, since the reconstruction process leverages spatial correlations across three orthogonal planes rather than operating independently on single slices, these noise-related effects exhibit spatial characteristics, which is commonly observed as not only as structured artifacts but also as degraded inter-slice continuity.

In summary, our primary goal is to train a generator *G* that transforms 𝒳 toward the clean domain, such that the resulting prediction 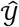 closely approximates 𝒴.

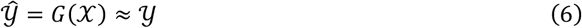

### 2.2. Network Architecture

In this study, we propose a novel generative adversarial denoising framework based on a dual-generator and dual-discriminator GAN architecture, enhanced by the integration of a Pseudo-Warping Module to promote spatial continuity. The overall architecture of our model is illustrated in **Fig. 1**(a), which comprises the following three key components:

**Fig. 1.**
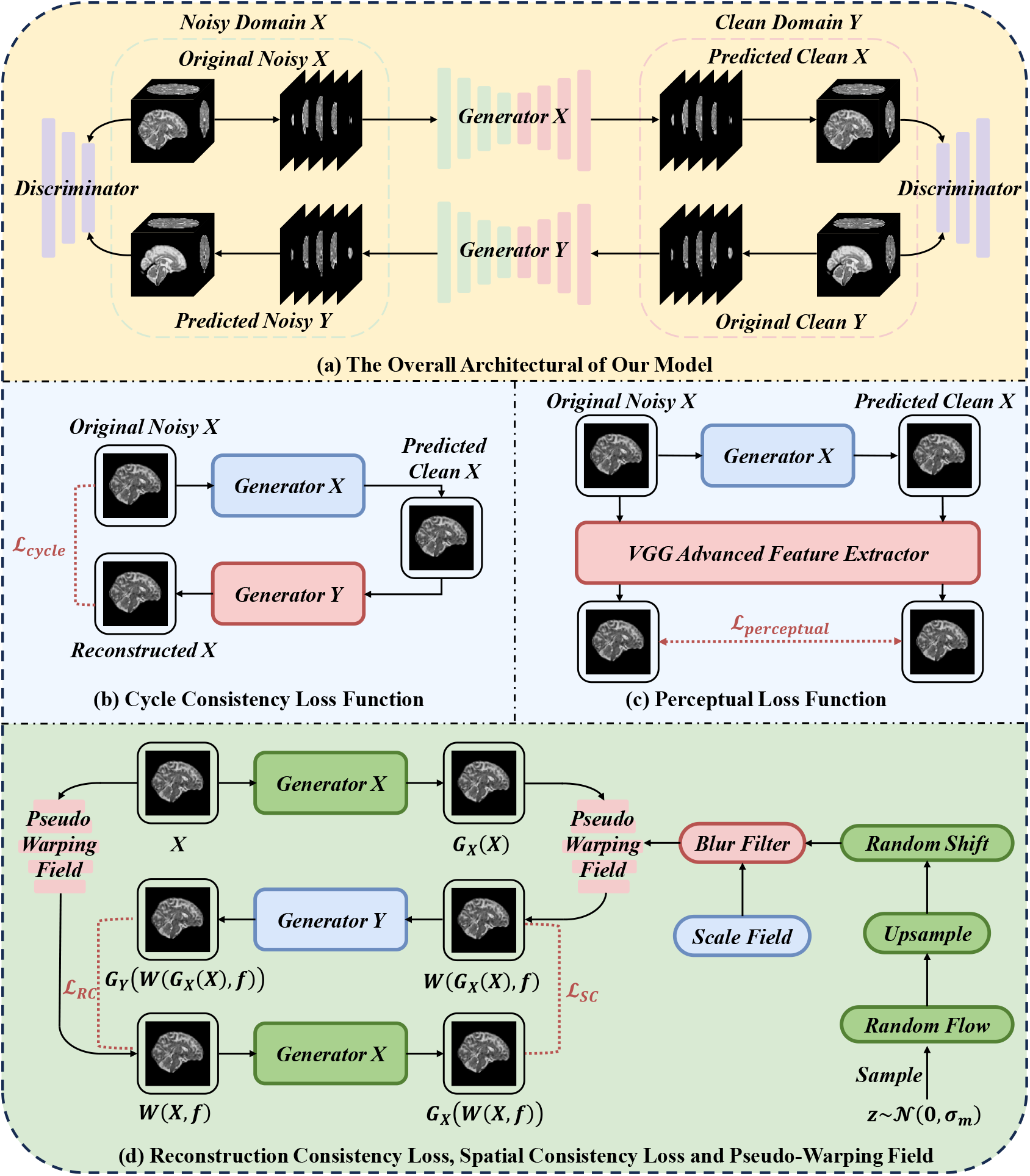
Overview of the proposed PW-GAN denoising framework. a) The Overall Architectural of Our Model. b) Cycle Consistency Loss Function. c) Perceptual Loss Function. d) Reconstruction Consistency Loss and Spatial Consistency Function

#### Generators

*G*_*X*_: Maps images from the noisy domain 𝒳 (thick-slice reconstructed MRI) to the clean domain 𝒴 (thin-slice reconstructed MRI).

*G*_*Y*_: Performs the inverse mapping from the clean domain 𝒴 back to the noisy domain 𝒳.

#### Discriminators

*D*_*X*_: Aims to distinguish between real noisy-domain images *X*and those synthesized by *G*_*X*_(*X*).

*D*_*Y*_: Discriminates between real clean-domain images *Y* and generated outputs *G*_*Y*_(*Y*).

#### Pseudo-Warping Field

*W*(*X, f*): Applies a deformation to the input image *X* guided by an generated optical flow field *f*, yielding a pseudo-slice that serves to improve inter-slice consistency.

It is worth noting that we formulate the denoising process as a cross-domain style transfer task, where the data from domains 𝒳 and 𝒴 are inherently unpaired. To ensure that the generators learn to suppress only noise-related features without altering underlying anatomical structures, we employ a carefully designed combination of loss functions. Furthermore, the incorporation of the pseudo-warping field renders our model more sensitive to inter-slice discontinuities and interpolation artifacts. This, in turn, endows the framework with an enhanced capacity to preserve and reinforce anatomical continuity across adjacent slices.

### 2.3. Pseudo-Warping Field

The pseudo-warping field consists of two components: a scale field and a shift field. The scale field applies subtle isotropic scaling to the input image, intended to simulate minor anatomical size variations across adjacent slices. The shift field, on the other hand, is constructed by first sampling a random optical flow from a standard normal distribution *z*∼𝒩(0, *σ*_*m*_), which is then upsampled to match the resolution of the original image. This step enhances the model’s sensitivity to local anatomical misalignments, such as those introduced by interpolation artifacts, and facilitates their correction. Subsequently, a slight random displacement is applied to the shift field, encouraging the network to better capture inter-slice continuity. Collectively, these operations help reinforce the consistency of 3D anatomical structures in the denoised outputs.

In conjunction with the pseudo-warping field, two additional consistency losses are employed—namely, the spatial consistency loss and the reconstruction cycle-consistency loss. The specific formulations and underlying mechanisms of these loss terms are detailed in the *Loss Function* section. Briefly, these two objectives work in concert to ensure that the final output remains consistent regardless of the order in which the generation operation *G* and the warping operation *W* are applied. This design encourages the network to develop a robust understanding of anatomical coherence under various spatial transformations.

### 2.4. Loss Functions

#### Adversarial Loss

The adversarial component, as shown in **Fig. 1**(a), is optimized through a dynamic minimax game between the generator and the discriminator. The generator is trained to minimize the adversarial loss by producing denoised images that closely resemble those from the target domain distribution, thereby aiming to deceive the discriminator. Conversely, the discriminator seeks to maximize its ability to distinguish between real and synthesized samples, thus driving the generator to continually improve the realism and fidelity of its outputs.

To improve training stability and enhance the visual fidelity of generated images, we adopt the least squares GAN (LSGAN) objective [42], which replaces the conventional binary cross-entropy loss with a least-squares regression formulation. Specifically, to encourage realistic image translation in both directions, the generator *G*_*X*_ maps noisy inputs from domain 𝒳 to their denoised counterparts in domain 𝒴, while *G*_*Y*_ performs the reverse mapping. The discriminators *D*_*X*_ and *D*_*Y*_ are trained to distinguish between real and synthesized images in their respective domains. The overall adversarial loss is defined as:

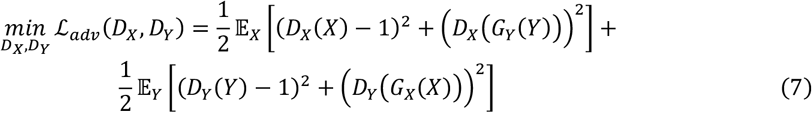

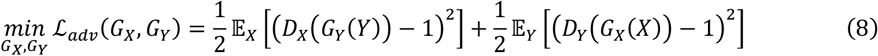

#### Cycle Consistency Loss

As illustrated in **Fig. 1**(b), to preserve the reversibility of cross-domain mappings and to mitigate the risk of anatomical information loss due to the generative diversity inherent in GANs, we incorporate a cycle consistency constraint. Specifically, the original noisy image *X* is first translated to the clean domain via the generator *G*_*X*_, yielding *G*_*X*_(*X*). This generated image is then mapped back to the noisy domain using the generator *G*_*Y*_. An L1 loss is computed between the reconstructed image *G*_*Y*_(*G*_*X*_(*X*)) and the original input *X*, encouraging the network to maintain structural fidelity throughout the transformation cycle.

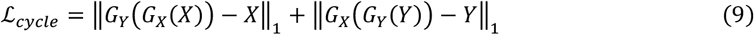

#### Perceptual Loss

As illustrated in **Fig. 1**(c), we employ a pretrained VGG network to extract advanced feature representations from both the original image *X* and its denoised counterpart *G*_*X*_(*X*). The mean squared error (MSE) between these feature maps is then computed to form the perceptual loss. By introducing constraints in the feature space, this loss encourages the preservation of fine anatomical details and structural integrity, ultimately enhancing the perceptual quality of the output as perceived by the human visual system.

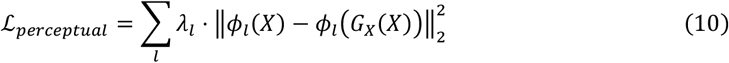

#### Consistency Loss

As introduced in Section 2.3, the consistency losses are designed to operate in conjunction with the pseudo-warping field, with the shared objective of suppressing interpolation artifacts and enhancing the continuity of 3D anatomical structures across slices. The overall mechanism is illustrated in **Fig. 1**(d).

#### Reconstruction Cycle-Consistency Loss

On one path, a noisy input image from domain 𝒳 is first translated into a denoised image via the generator *G*_*X*_(*X*). A pseudo-warping field *f* is then derived based on this output and applied, resulting in a warped image *W*(*G*_*X*_(*X*), *f*). This warped image is subsequently mapped back to the noisy domain 𝒳 using the generator *G*_*Y*_, yielding *G*_*Y*_(*W*(*G*_*X*_(*X*), *f*)). In parallel, we directly apply the same warping field *f* to the original noisy input *X*, obtaining *W*(*X, f*). The reconstruction cycle-consistency loss is then computed by comparing *W*(*X, f*) and *G*_*Y*_(*W*(*G*_*X*_(*X*), *f*)).

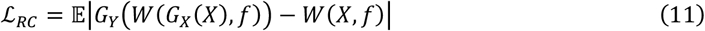

#### Spatial Consistency Loss

Building upon the warped image *W*(*X, f*), we further transform it into the clean domain 𝒴 using *G*_*X*_, resulting in *G*_*X*_(*W*(*X, f*)).Then, we compute a spatial consistency loss by comparing it against *W*(*G*_*X*_(*X*), *f*).

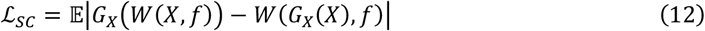

Through the combined effect of these two consistency losses, the framework is encouraged to produce anatomically coherent outputs, invariant to the order in which generation *G* and warping *W* operations are applied.

In summary, the overall loss function of the proposed model *ℒ* can be formulated as follows:

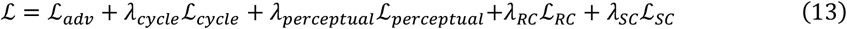

### 2.5. Evaluation Metrics

To enable a more comprehensive evaluation of the model’s performance, we incorporate not only qualitative assessments but also a set of quantitative metrics to measure denoising effectiveness. Specifically, Peak Signal-to-Noise Ratio (PSNR), Structural Similarity Index Measure (SSIM), and Root Mean Squared Error (RMSE) are employed to evaluate both image fidelity and the preservation of anatomical structures. It is worth noting, however, that these quantitative metrics require access to ground truth references for valid comparison. Consequently, in experiments involving real clinical data—where such ground truth is typically unavailable—quantitative evaluation cannot be reliably conducted.

In addition, to assess the inter-slice continuity in the denoised fetal brain MRI volumes, we introduce a metric termed Warping Error (WE) [43]. This metric combines L1 norm with an exponentially weighted L2 norm to provide a more nuanced assessment of anatomical consistency across adjacent slices. Intuitively, a lower WE value indicates higher spatial coherence and improved preservation of anatomical continuity in three dimensions.

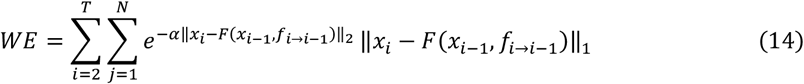

where *T* denotes the number of adjacent slices in the same direction from a single fetal brain MRI volume, and *N* represents the total number of pixels in a single slice.

## 3. Experiments

### 3.1. Datasets and Experiments Setup

In this study, we utilized a clinical dataset comprising fetal brain T2-weighted MR images collected from 228 cases at the Shenzhen Maternity & Child Healthcare Hospital. The imaging was performed using a 1.5-Tesla Philips MRI scanner, with gestational ages ranging from 22 to 39 weeks (mean ± standard deviation: 33.7 ± 3.8 weeks). For each subject, at least three orthogonal stacks were acquired in coronal, sagittal, and axial orientations. These stacks were anisotropic, with an in-plane resolution of 0.8 mm and a slice thickness of 5 mm. The acquisition parameters included a repetition time (TR) of 1.1 seconds, an echo time (TE) of 100 milliseconds, and a flip angle of 90 degrees.

To ensure the integrity of the dataset, subjects with suspected brain developmental abnormalities were excluded. Subsequently, the clinical scans were processed using NeSVoR [44] to perform super-resolution reconstruction, resulting in isotropic volumes of 256×256×256 voxels with a final slice thickness of 0.8 mm.

Additionally, we incorporated the publicly available CRL Atlas dataset [40] as a reference for clean-domain images. This dataset was constructed from MRI scans of 81 normally developing fetuses, covering a gestational age range from 21 to 38 weeks. The dataset provides super-resolved reconstructions at a resolution of 256×256×256 voxels, which are readily available for reference purposes in our study.

To ensure anatomical consistency across developmental stages, the clinical samples were first grouped according to gestational age and matched with corresponding age-appropriate, noise-free templates from the Spatial alignment between the clinical data and the CRL templates was verified using 3D Slicer, confirming conformity to the RAS coordinate system and correcting for any initial positional or orientational discrepancies. Rigid registration of the clinical volumes to the atlas space was then performed using the FSL-FLIRT tool with a 12-degree-of-freedom affine transformation. Subsequently, orthogonal slicing was applied along the sagittal, coronal, and axial planes, with slices lacking brain tissue entirely being excluded from further experiments.

For robust model evaluation, we adopted a five-fold cross-validation strategy. 80% of the dataset was allocated for training and validation, while the remaining 20% was reserved for independent testing.

### 3.2. Clinical Data Denoising Experiments

To assess the applicability of our model in real-world clinical scenarios—where paired noisy-clean image data are typically unavailable—we conducted denoising experiments on clinical fetal brain MRI data. Following the preprocessing and dataset partitioning procedures described in Section 3.1, we designated the high-resolution CRL Atlas reconstructions as clean reference images (i.e. the clean domain), and treated the clinical reconstructions as noisy inputs (i.e. the noisy domain). The denoised outputs produced by our model were then qualitatively compared against several widely adopted unsupervised baselines, including Deformed2Self, Noise2Void and OTE-GAN.

It is worth noting that supervised approaches such as DnCNN were not applicable in this context due to their dependence on explicitly paired noisy and clean data, which are unavailable in our setting. Although Noise2Noise is categorized as an unsupervised method, it requires multiple independent noisy observations of the same target—an assumption that is not satisfied in the single-slice acquisition paradigm of routine clinical practice. As such, it was also excluded from comparison.

### 3.3. Simulated Data Denoising Experiments

To enable a more comprehensive evaluation of our model’s performance—both qualitatively and quantitatively—against existing supervised and unsupervised denoising and optimizing methods, we further conducted experiments on a synthetic dataset with simulated noises constructed from the CRL Atlas. Specifically, we adopted the CRL Atlas dataset as the ground truth and generated corresponding noisy images by simulating realistic MRI artifacts based on the noise formulation introduced in Section 2.1. The simulated noise is designed to replicate artifacts commonly encountered in real scanning scenarios, comprising three components in our experiments: periodic artifacts, inter-slice discontinuity artifacts, and system-level random noise. Following Equation (5), periodic artifacts and local signal mixing noise are introduced to mimic the effects of residual noise in thick-slice reconstruction process, denoted as ℛ(*n*). We additionally simulated inter-slice discontinuities arising during reconstruction 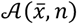 by incorporating staircase artifacts, and added random noise to account for potential intrinsic system-level perturbations *ϵ*. This procedure yielded a paired noisy-clean dataset that served as the basis for our further denoising experiments. An example of the resulting image pairs is illustrated in **Fig. 2**.

**Fig. 2.**
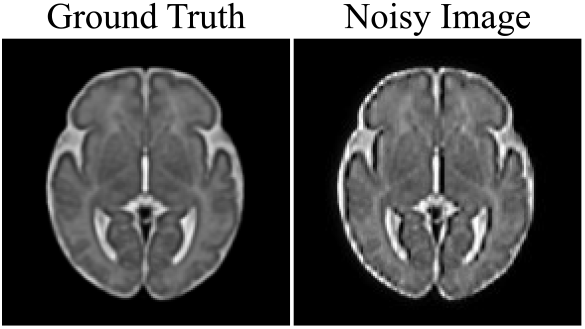
Noisy Images Simulated via Noise Modeling and Corresponding Ground Truth

During evaluation, in addition to the aforementioned unsupervised baselines, we included two supervised methods—DnCNN and Noise2Noise for comparison. Multiple criteria were employed, including visual inspection and widely used quantitative metrics such as PSNR, SSIM, RMSE. Furthermore, to evaluate the improvement in three-dimensional anatomical continuity before and after denoising, we utilized the warping error (WE) metric described in Section 2.5 for evaluation. Notably, although our model used the CRL Atlas data as clean references during this experiment, the data employed for training our model remained unpaired, thus preserving the integrity of the unsupervised training paradigm.

### 3.4. Generalization Ability Evaluation

To evaluate the denoising performance and generalization capability of our model under different noise conditions, we conducted additional experiments on a separate set of simulated data. Specifically, we introduced Rician noise with a standard deviation of 15% and salt-and-pepper noise at a 5% level to the CRL Atlas dataset, thereby simulating challenging noise scenarios. **Fig. 3** presents representative examples of simulated noisy images alongside their corresponding ground truth. The corrupted images were then processed using our model, and the outputs were compared—both visually and quantitatively— with those produced by representative supervised and unsupervised denoising approaches. The results of this evaluation are presented in Section 4.3.

**Fig. 3.**
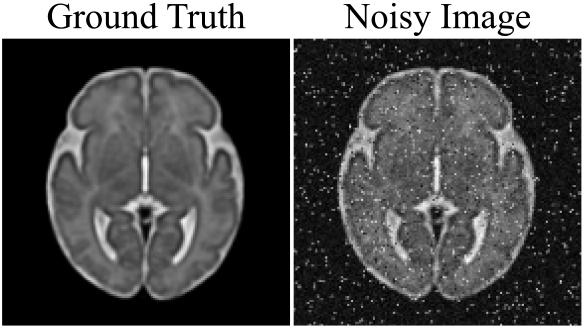
Ground Truth and Simulated Noisy Image Corrupted with Rician Noise (*σ*=15%) and Salt-and-Pepper Noise (5%)

### 3.5. Ablation Studies

To investigate the contribution of the pseudo-warping field and associated loss functions to the denoising and optimization performance of our model, we conducted a quantitative evaluation of these components within the context of the synthetic denoising experiments. Specifically, we compared the outcomes under various model configurations to assess their respective impacts on denoising quality and training stability.

### 3.6. Implementation Details

During the training phase, the model was optimized using an *NVIDIA A800 GPU* equipped with 80*GB* of memory. The batch size was set to 22. Several loss components were incorporated into the objective function with the following weights: *λ*_*cycle*_ = 20.0, *λ*_*perceptual*_= 1.0, *λ*_*RC*_ =10.0, *λ*_*SC*_=10.0. Training was conducted over 50 epochs, with the learning rate initialized at 5 × 10^−5^ for the first 30 epochs. In the subsequent 20 epochs, the learning rate was gradually decayed to zero to facilitate convergence.

## 4. Results

### 4.1. Clinical Data Denoising Result

**Fig. 4** illustrates a comparison between our method and existing unsupervised denoising approaches on clinical fetal brain MRI images. As shown, the original scans exhibit substantial noise interference, which not only diminishes the visual clarity of neuroanatomical structures but may also hinder the accuracy and robustness of downstream tasks such as fetal brain tissue segmentation.

**Fig. 4.**
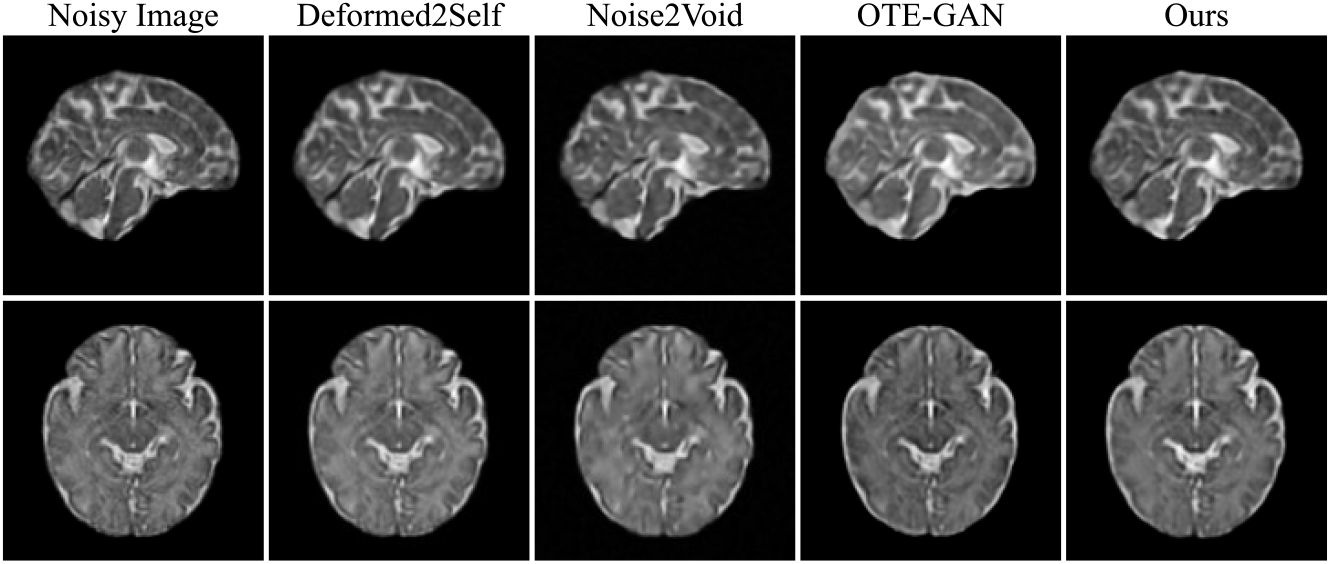
Visualization results of different unsupervised methods on real clinical fetal brain MRI data

Our model demonstrates competitive, and in some cases superior, performance while maintaining strong anatomical consistency. Notably, thanks to the ability of our framework to incorporate a designated reference dataset as clean guidance, it produces more realistic outputs without introducing excessive smoothing artifacts often observed in other methods.

### 4.2. Simulated Data Denoising Result

We compared the proposed method with several state-of-the-art supervised and unsupervised denoising approaches on the dataset described in Section 3.3. **Fig. 5** presents the qualitative evaluation of denoising performance across different methods. To better illustrate the differences among the results, we generated pixel-wise difference maps for each denoised output. These maps were computed by first normalizing the outputs of all models and then calculating the absolute differences between each denoised image and its corresponding ground truth.

**Fig. 5.**
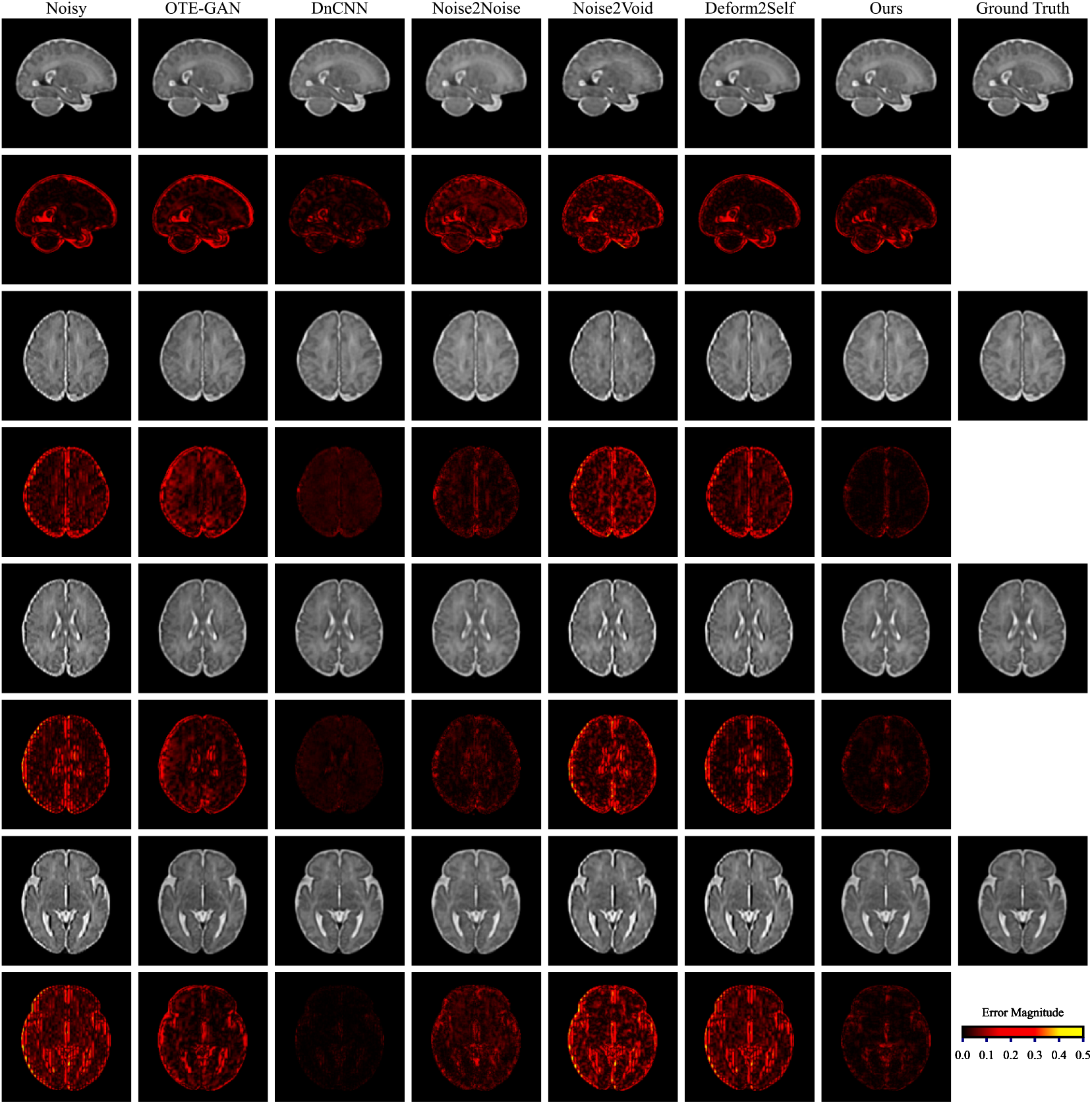
Visualization results of different methods on the CRL Atlas dataset with simulated noise. For each group of images, a difference map with respect to the ground truth is also provided, where darker regions in the difference maps reflect closer similarity between the denoised result and the ground truth.

In the resulting visualizations, regions with larger discrepancies are highlighted, indicating areas where anatomical structures may have been altered during denoising. Conversely, areas with minimal differences appear darker, with black representing perfect agreement with the ground truth.

As illustrated in **Fig. 5**, the original noisy images exhibit noticeable degradation in visual quality, primarily due to interpolation artifacts and discontinuities in anatomical structures introduced during the reconstruction process. After denoising, our method achieves the best visual performance among all unsupervised approaches and delivers results that are comparable to, or in some cases surpass, those of supervised methods such as DnCNN. Notably, in restoring fine anatomical details, our model is particularly effective at eliminating artifact-related interference and reconstructing structurally coherent and visually sharp images.

To further validate the denoising performance, we conducted a quantitative evaluation using standard metrics including PSNR, SSIM, and RMSE, as presented in **Table 1**.

**Table 1.** Quantitative evaluation of denoising performance on the CRL Atlas dataset with simulated noise. All metrics are computed using the original noise-free paired CRL Atlas data as ground truth. The top two results for each metric are highlighted in bold.

| Method | PSNR(↑) | SSIM(↑) | RMSE(↓) |
| --- | --- | --- | --- |
| DnCNN | **46.34** | **0.9975** | **0.0057** |
| OTE-GAN | 34.65 | 0.9854 | 0.0198 |
| Deformed2Self | 27.07 | 0.9329 | 0.0564 |
| Noise2Noise | 28.41 | 0.9100 | 0.0381 |
| Noise2Void | 31.24 | 0.9446 | 0.0291 |
| Ours | **39.65** | **0.9932** | **0.0126** |

Although DnCNN achieved the highest scores in both qualitative and quantitative assessments, it relies on paired clean-noisy image data for training. In contrast, our method produces highly competitive results without requiring such paired data.

Furthermore, as observed in **Fig. 5**, our model demonstrates superior capability in mitigating interpolation artifacts and anatomical discontinuities introduced during reconstruction. This advantage can be attributed to the incorporation of pseudo-warping fields and two consistency losses, which enable the model to perceive and correct misalignments across adjacent slices. **Table 2** further supports this observation by reporting the warping errors of denoised results across different anatomical planes, where our method consistently shows improved structural continuity.

**Table 2.** Wrapping errors of different denoising models evaluated across multiple anatomical planes. The top one result for each plane is highlighted in bold.

| Method | Warping Error ($\times 10^{-3}$) ↓ | | |
| --- | --- | --- | --- |
|  | Coronal Axis | Transverse Axis | Sagittal Axis |
| DnCNN | **10.13** | 8.62 | 7.54 |
| OTE-GAN | 11.18 | 9.33 | 8.27 |
| Deformed2Self | 15.43 | 12.10 | 10.81 |
| Noise2Noise | 12.56 | 8.76 | 7.81 |
| Noise2Void | 17.56 | 13.59 | 12.42 |
| Ours | 11.28 | **8.47** | **7.12** |

In summary, our model effectively removes residual noise and interpolation artifacts from fetal brain MRI after reconstruction and significantly enhances the continuity of anatomical structures, all without relying on paired clean-noisy image pairs.

### 4.3. Generalization Ability Evaluation Results

As illustrated in Section 3.4, we employed a combination of Rician noise (*σ*=15%) and salt-and-pepper noise (5%) to evaluate the generalization capability of our model under alternative noise conditions. A visual comparison of the denoising results is presented in **Fig. 6**. As detailed, unsupervised models trained solely on noisy images struggle to recover clean structures in such severely degraded scenarios, largely due to the absence of clean image guidance. In contrast, our model benefits from learning the distribution of clean-domain images, enabling it to generate significantly clearer results under the same conditions. While DnCNN still yields the visually cleanest results, it relies on paired data for training. In this regard, our approach offers a practical advantage in clinical applications, where acquiring paired noisy-clean datasets is often infeasible.

**Fig. 6.**
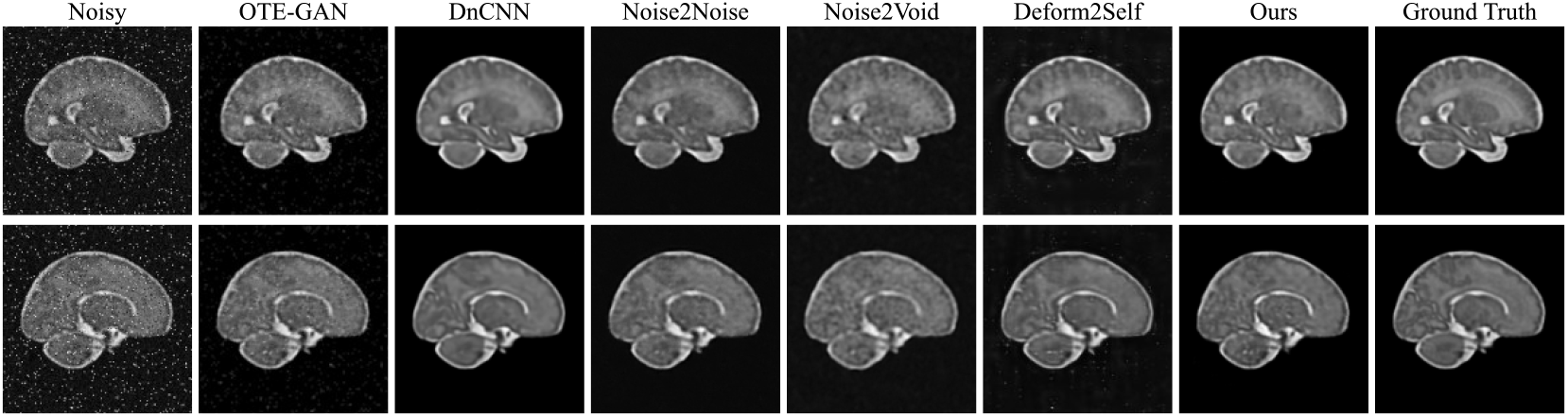
Visual Comparison of Denoising Results on Simulated Noisy Images Corrupted with Rician Noise (*σ*=15%) and Salt-and-Pepper Noise (5%)

Quantitative evaluations using PSNR, SSIM, and RMSE are summarized in **Table 3**. From a numerical perspective, our method achieves performance on par with supervised approaches and surpasses other unsupervised baselines. Additionally,

**Table 3.** Quantitative Evaluation of the Model on Simulated Noisy Images Corrupted with Rician Noise (*σ*=15%) and Salt-and-Pepper Noise (5%). The top two results for each metric are highlighted in bold.

| Method | PSNR(↑) | SSIM(↑) | RMSE(↓) |
| --- | --- | --- | --- |
| DnCNN | **44.33** | **0.9986** | **0.0065** |
| OTE-GAN | 29.57 | 0.8902 | 0.0333 |
| Deformed2Self | 21.37 | 0.7051 | 0.0855 |
| Noise2Noise | 28.54 | 0.8138 | 0.0374 |
| Noise2Void | 25.89 | 0.7711 | 0.0515 |
| Ours | **37.19** | **0.9912** | **0.0148** |

**Table 4** reports the results of warping error analysis. Benefiting from the combined effect of the pseudo-warping field and consistency loss, our model substantially improves anatomical continuity across slices.

**Table 4.**
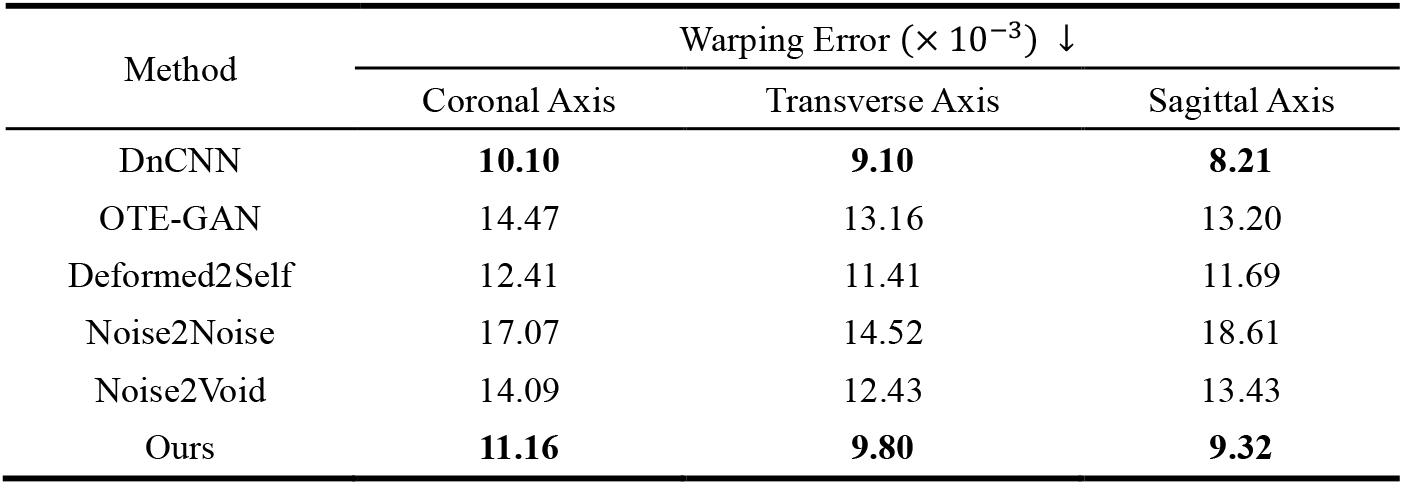
Wrapping errors of different denoising models evaluated across multiple anatomical planes. The top two results for each metric are highlighted in bold.

| Method | Warping Error ($\times 10^{-3}$) ↓ | | |
| --- | --- | --- | --- |
|  | Coronal Axis | Transverse Axis | Sagittal Axis |
| DnCNN | **10.10** | **9.10** | **8.21** |
| OTE-GAN | 14.47 | 13.16 | 13.20 |
| Deformed2Self | 12.41 | 11.41 | 11.69 |
| Noise2Noise | 17.07 | 14.52 | 18.61 |
| Noise2Void | 14.09 | 12.43 | 13.43 |
| Ours | **11.16** | **9.80** | **9.32** |

Interestingly, this experiment also revealed an unexpected advantage of treating denoising as a form of style transfer: the model becomes more aware of the contextual relationship between anatomical regions and the background. This perspective proves especially beneficial for post-reconstruction fetal brain MRI denoising, where background noise often interferes with image quality. By leveraging this implicit structural understanding, our model demonstrates enhanced capability in suppressing background noise and preserving anatomical fidelity.

### 4.4. Ablation Study Results

In the ablation study, we individually removed the RC loss, SC loss, pseudo-warping field, and VGG loss to investigate the contribution of each component to image denoising performance and the enhancement of anatomical consistency. The quantitative results are summarized in **Table 5** and **Table 6**.

As shown in the results, removing the RC loss, SC loss, or pseudo-warping field leads to a noticeable decline in both denoising quality and spatial anatomical consistency. In contrast, the exclusion of the VGG loss primarily affects denoising performance, with relatively minor impact on anatomical continuity across slices. These findings suggest that the incorporation of the pseudo-deformation field and consistency losses plays a pivotal role in preserving structural coherence across adjacent slices.

**Table 5.** Quantitative performance of our model after ablating individual component. The most influential module for each metric is highlighted in bold.

| Module | PSNR(↑) | SSIM(↑) | RMSE(↓) |
| --- | --- | --- | --- |
| Base Line | 39.65 | 0.9932 | 0.0126 |
| w/o RC Loss | 34.96 | **0.9867** | 0.0191 |
| w/o SC Loss | **34.85** | 0.9877 | **0.0198** |
| w/o Warping Field | 36.01 | 0.9891 | 0.0174 |
| w/o VGG Loss | 37.15 | 0.9906 | 0.0159 |

**Table 6.** Warping errors across different anatomical planes after ablating individual component of our model. The most influential module for each metric is highlighted in bold. This metric reflects the contribution of each component to enhancing spatial anatomical consistency.

| Method | Warping Error ($\times 10^{-3}$) ↓ | | |
| --- | --- | --- | --- |
|  | Coronal Axis | Transverse Axis | Sagittal Axis |
| Base Line | 11.28 | 8.47 | 7.12 |
| w/o RC Loss | 12.75 | 9.23 | 8.62 |
| w/o SC Loss | **13.80** | **9.46** | **8.98** |
| w/o Warping Field | 13.56 | 8.61 | 7.81 |
| w/o VGG Loss | 11.98 | 8.65 | 7.21 |

## 5. Conclusions

In this study, we propose a novel framework for post-reconstruction denoising and structural refinement of fetal brain MRI, based on pseudo-warping field. By formulating the task as an inter-domain translation problem between noisy and clean domains, our method is capable of reducing residual noise and artifacts, while also reinforcing anatomical continuity—without the need for paired training data.

Traditional unsupervised methods, lacking access to clean reference images, often struggle to correct structural distortions and interpolation artifacts. Meanwhile, supervised approaches, though typically more effective, depend heavily on paired clean images, which are often unavailable in clinical practice and thus limit their real-world applicability. By introducing unpaired clean images as guidance through inter-domain translation, our method offers a balanced alternative between supervised and unsupervised paradigms—achieving strong denoising performance without the reliance on clean-noisy pairs. Furthermore, the integration of a VGG-based perceptual module enhances the model’s ability to preserve high-level structural features such as textures and edges. Most importantly, the use of pseudo-warping field combined with consistency losses equips the model with a refined sensitivity to interpolation artifacts and anatomical discontinuities. This enables the generation of denoised results that are not only high in visual fidelity, but also markedly improved in terms of spatial anatomical consistency across 3D reconstructions.

